# Using the Price equation to analyze multi-level selection on the reproductive policing mechanism of bacterial plasmids

**DOI:** 10.1101/079574

**Authors:** Kyriakos Kentzoglanakis, Sam P. Brown, Richard A. Goldstein

## Abstract

The replication control system of non-conjugative bacterial plasmids constitutes a simple and elegant example of a reproductive policing mechanism that moderates competition in the intra-cellular replication pool and establishes a mutually beneficial partnership among plasmids within a bacterial host and between plasmids and their hosts. The emergence of these partnerships is a product of the conflict between the evolutionary interests of hosts, who seek to maximize their growth rates within the population, and plasmids, who seek to maximize their growth rates within hosts. We employ a multi-scale computational model describing the growth, division and death of hosts, as well as the independent replication of plasmids within hosts, in order to investigate the implications of this conflict for the evolution of the plasmid replication parameters. We apply the multi-level form of the Price equation in order to quantify and elucidate the various selective pressures that drive the evolution of plasmid replication control. Our analysis shows how the evolution of the constituent components of the plasmid replication control system are shaped by selection acting at the level of hosts and the level of plasmids. In addition, we calculate finer-grained selective pressures that are attributed to atomic plasmid-related events (such as intra-cellular replication and plasmid loss due to host death) and demonstrate their special role at the early stages of the evolution of policing. Our approach constitutes a novel application of the Price equation for discerning and discussing the synergies between the levels of selection given the availability of a mechanistic model for the generation of the system’s dynamics. We show how the Price equation, particularly in its multi-level form, can provide significant insight by quantifying the relative importance of the various selective forces that shape the evolution of policing in bacterial plasmids.

## Introduction

The emergence and maintenance of cooperation has always been a challenging problem in evolutionary biology, primarily characterized by the stark conflict between an individual’s pursuit of immediate reproductive gains, achieved through competing with other individuals within a collective, and the success of the collective, achieved through cooperative behaviour and consequently enjoyed by all individuals therein. Competition between individuals within a collective for access to fitness-increasing resources can lead to the “tragedy of the commons” [1], in which the over-exploitation and eventual exhaustion of such resources cause the population’s collapse. The tragedy of the commons can be averted by the moderation of individual competitiveness through social interaction mechanisms that promote cooperation. For example, in situations where there is high genetic relatedness among the individuals involved, kin selection can act in favor of the evolution of self-restraint through the carriage and transmission of shared genes that encode cooperative behaviour [2]. At low genetic relatedness, the maintenance of costly cooperation can be achieved through individual investment in policing mechanisms that repress competition between group members, thus increasing the group’s efficiency and productivity [3–6].

A simple and elegant example of reproductive policing for cooperation between species is the replication control system of bacterial plasmids. Bacterial plasmids are extra-chromosomal genetic elements that replicate autonomously within bacterial cells. They are organized as collections of discrete genetic modules [7, 8] which encode functions necessary for their maintenance and propagation within bacterial populations. These functions include replication control and horizontal (conjugative) transfer, as well as systems that minimize the probability of segregational loss such as multimer resolution [9], active partitioning during cell division [10] and toxin-antitoxin complexes [11].

The plasmid-coded replication control system is a mechanism for repressing competition between plasmids in the intra-cellular replication pool. The system works by regulating plasmid replication so as to fix a “characteristic” copy number and restrict variation around that number. Plasmid copy number control (CNC) is implemented by means of a negative feedback loop that responds to copy number fluctuations by appropriately modulating individual plasmid replication rates [12–14]. The CNC mechanism is a policing system characterized, first, by the production of rate-limiting constitutive factors such as Rep proteins [15, 16] or RNA primers [17] that act in cis and are responsible for the initiation of plasmid replication (individual competitiveness/selfishness); second, by the synthesis of trans-acting replication inhibitors (policing resources) that act upon the target initiation factors [18]; and, third, by the binding of replication inhibitors to plasmid-specific sites (obedience to policing).

The cis- and trans- acting elements of the CNC system are subject to a dynamic evolutionary conflict between two levels of selection [19, 20]: at the intra-cellular level, cis-selfish mutations [20] that increase the rate of initiation of plasmid replication or reduce the binding affinity to the trans-acting inhibitor would result in plasmids that proliferate faster in comparison to more prudent plasmids with a higher degree of adherence to the CNC mechanism. Hence, within-host selection will favor cis-selfishness, thus exacerbating intra-cellular competition between plasmids. However, hosts bearing increasingly selfish and disobedient plasmids will grow slower compared to fellow hosts with stricter replication control among plasmids. This way, selective pressures towards cis-selfishness at the intra-cellular level are counter-balanced by selection at the level of hosts that penalizes hosts with selfish plasmids and, therefore, selfish plasmids themselves.

A powerful conceptual framework for understanding the evolution of various traits is the Price equation, a mathematical identity that relates selection to covariance [21]. This equation has been adapted to the situation of multi-level selection where there is conflicting selection acting at the individual and collective level [26, 27], such as in the case of policing. There are, however, two significant barriers that limit the application of the Price equation. Firstly, the various quantities described in the Price equation, such as the covariance of a trait with replicative fitness, are difficult to measure. Secondly, the Price equation represents the relationship between statistical measures such as averages and covariances, which are incomplete descriptions of the state of the system. As a result, the Price equation cannot be used to predict future behavior of a population without additional assumptions [31]. Neither of these aspects are a limitation to the use of the Price equation to analyse evolutionary simulations. Such simulations are completely transparent – the state of every individual at every time point is available to the researcher with no uncertainty. In addition, the Price equation is not needed for generating the dynamics, only *a posteriori* analysis of the evolutionary process.

In this paper, we use a previously described computational agent-based model [22] that characterizes the mechanics of host growth and autonomous plasmid replication in order to simulate the evolution of the components of the plasmid replication control system. We then use the Price equation to quantify the effects of selection on the plasmid replication parameters, not only across the levels of selection but also across specific types of events (such as plasmid replication and loss due to host death) that determine plasmid fitness. We demonstrate the power of the multilevel Price equation to analyse such evolutionary simulations, enabling us to analyze and describe the synergies that shape the evolution of reproductive policing in bacterial plasmids.

## Methods

### Model

We use a previously described model [22] that defines the mechanics of the asynchronous growth, division and death of hosts in a bacterial population, as well as the independent replication of plasmids within hosts. More specifically, we consider a population of hosts which grow and divide or die and impose a fixed maximum population capacity (*N* = 1000) so that, when the population operates below that capacity, daughter cells are added to the growing population. When the population operates at maximum capacity, the two daughter cells resulting from a cell division event replace the parent cell and a randomly selected cell from the population so that, at maximum capacity, every birth (cell division) corresponds to a random death. We assume that plasmids do not encode an explicit partitioning mechanism, so that when a cell divides, its plasmids are distributed among daughter cells on the basis of binomial segregation (i.e. each plasmid is equally likely to end up in either of the daughter cells).

Our model for cellular growth assumes that cell metabolism produces biomass and that each host *i* grows independently by updating its biomass Ω_*i*_. When Ω_*i*_ ≥ 2Ω_0_ (with Ω_0_ = 1), host *i* divides and its plasmids are distributed among the two daughter cells, while when Ω_*i*_ < 0 the host dies as a result of a negative balance between metabolic benefits and costs. The biomass Ω_*i*_ of each host is updated at each time step of the simulation according to:

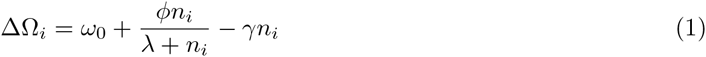

where *n*_*i*_ is the total number of plasmid copies in the host and *ω*_0_ is the cell’s basal growth rate. Parameters *ϕ* and *λ* characterize the beneficial contribution of the plasmid trait to cell metabolism which saturates at large *n*, while *γ* specifies the cost of maintenance (including the costs of gene expression, replication etc.), which is proportional to the copy number [23–25]. By differentiating the right-hand side of Equation 1, setting to zero and solving for *n*, we find the copy number 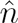 that is optimal for host growth given by:

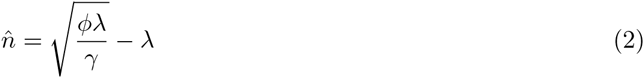

Within each host *i* in the population, plasmids replicate autonomously by competing in the intra-cellular replication pool. We assume that plasmids do not encode the neccessary functionality for mobilization and horizontal transmission across bacterial hosts, so that transmission occurs purely vertically from parent to daughter cells. Intra-cellular competition between plasmids is subject to regulation by the plasmid-coded CNC system, which operates on the basis of the production of plasmid-coded trans-acting replication inhibitors that bind to and deactivate a plasmid-specific target, thereby down-regulating the individual plasmid replication rate [18]. The replication rate of plasmid *j* in host *i* is given by:

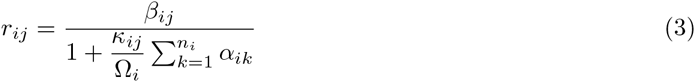

where *β*_*ij*_ is the plasmid’s basal replication rate and *κ*_*ij*_ is its binding affinity to the generic replication inhibitor, which is synthesized by each plasmid at a rate *α*_*ij*_/*τ*_*D*_ per plasmid, where *τ*_*D*_ is the (short) average lifetime of the inhibitor, resulting in a total inhibitor concentration of 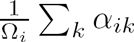 in host *i,* as the inhibitor is diluted with increasing host biomass Ω_*i*_. As the binding affinities are constrained by the limits of physico-chemical interactions, we set *κ* ∈ [0,1]. Similarly, we assume that the basal replication rate *β* and the rate *α* of inhibitor production are limited by physiological and biochemical constraints, therefore we also set *β, α* ∈ [0,1].

The replication of a plasmid implies the possibility of mutation with probability *µ*, in which case exactly one of its parameter values (*β*, *κ* or *α*, chosen at random) is modified. Let *v* ∈ [0,1] be the value of the parameter to be mutated. We choose the new parameter value *v′* from a uniform distribution 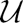 according to:

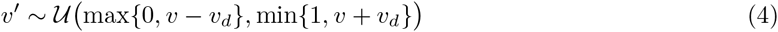

where *v_d_ =* 0.05 determines the maximum possible deviation in value that mutations are allowed to make.

In effect, our model partitions the plasmid replication control system into three distinct plasmid traits, consisting of each individual plasmid’s intrinsic level of selfishness *β,* its production of trans-acting policing resources *α* and its obedience *κ* to policing. The interaction between these components within each host determines the operation and efficiency of the CNC mechanism.

### Partitioning selection on plasmid replication parameters

Having defined the plasmid replication parameters, we are now interested in how they are influenced by the interplay between the fitness of the individual and the fitness of the collective that drives the emergence of stable replication control. The fitness of the collective is implicit in our simulations and depends on the rate of growth of individual hosts as a function of the plasmid copy number (local population size), with faster growth rates indicating shorter cell cycles and, therefore, more frequent division events (see Equation 1).

The fitness *f*_*ij*_ of an individual plasmid *j* in host *i* is the (actual) copy number of plasmid *j* in the population at the next time step of the simulation. This number includes the plasmid itself (if host *i* remains alive), any new copies and any possible losses as follows:

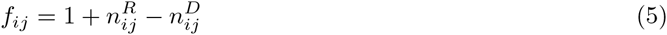

where 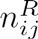 is the number of intra-cellular replication events and 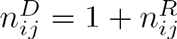 if host *i* dies (either due to ∆Ω < 0 or by coupling to the division of another host), otherwise 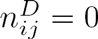. The latter implies that, in the event that host *i* dies, plasmid *j* disappears from the population.

The specification of a functional form for plasmid fitness motivates the consideration of the Price Equation as a means for investigating the selective pressures on the plasmid replication parameters. The Price Equation [21] is a mathematical identity that describes the change in the average value of a phenotypic character *z* (e.g. one of the plasmid replication parameters *β*, *κ*, or *α*) as a function of the statistical relationship (covariance) between fitness *f* and the character value *z* among individuals in a population, and the average fitness-weighted errors in the transmission of the character *z* from parent *i* to its offspring (transmission bias), according to:

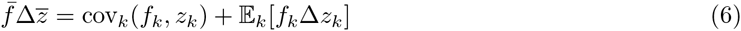

where 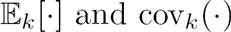 denote the expectation and covariance over all individuals *k* in the population respectively. In our case, this iteration would involve all plasmids in the population, irrespective of the host (collective) to which they belong. In essence, the covariance term in Equation 6 captures the portion of 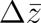 that is attributed to selection on *z,* while the expectation term captures the portion of 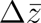 that is attributed to the fitness-weighted transmission errors ∆*z* during plasmid replication. The latter is expressed as the difference between each plasmid’s parameter value and the average value of its offspring, hence the transmission error for plasmid *k* with character value *z*_*k*_ can be expressed as:

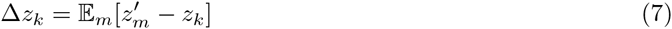

where *m* enumerates the *f*_*k*_ offspring of plasmid *k* at the next time step, each having a character value 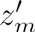.

The Price equation can also be rewritten to account for individuals that are nested within collectives [26, 27], in which case the covariance term representing selection on the phenotypic character *z* can be partitioned into two terms, corresponding to the portions of cov(*f*, *z*) that are attributed to selection between hosts and within hosts respectively, according to:

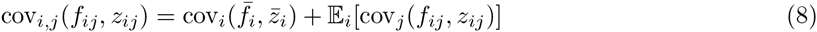

where *i* enumerates hosts, *j* enumerates plasmids in host *i* with plasmid copy number *n*_*i*_, and 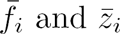 are the average fitness and character values of plasmids in host *i* respectively, with 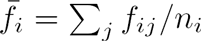 and 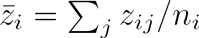.

The calculation of the covariance and expectation over *i* in Equation 8 should also take into account that hosts *i* differ with respect to the local population size *n*_*i*_ (plasmid copy number). For this reason, the contribution of each host to the expectation and covariance terms should be weighted by the host’s copy number [26, 27], according to:

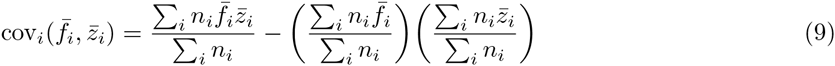

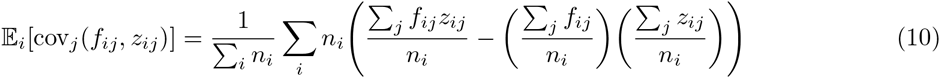

By substituting Equation 8 into Equation 6, we obtain the multi-level version of the Price equation:

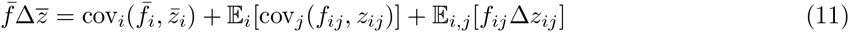

in which the first term captures selection on plasmid collectives at the level of hosts (between-host selection), the second term captures selection on individual plasmids at the intra-cellular level (within-host selection), while the third term quantifies the bias in the transmission of character *z* during plasmid replication.

### Attributing selection to specific types of events

The components of Equation 11 can be further partitioned into the constituent components of plasmid fitness. This can be achieved by substituting Equation 5 into the covariance term of the Price equation:

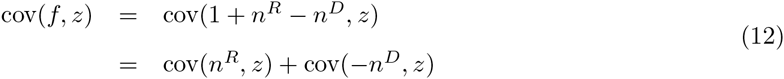

where selection on character *z* is partitioned into two components corresponding to the classes of events that affect the survival and propagation of plasmids, namely intra-cellular replication (*n*^*R*^) and possible losses of plasmid copies due to cellular death (*n*^*D*^). As such, between-host selection on character *z* can be decomposed according to:

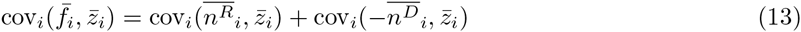

where 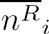 is the average number of intra-cellular replication events within host *i* and 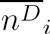 is the average number of plasmid losses if host *i* dies.

Similarly, within-host selection on character *z* can be decomposed according to:

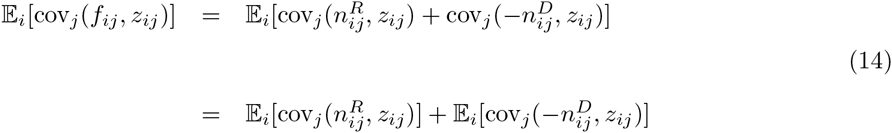

In effect, each component in these equations captures the portion of selection on character *z* that can be attributed to changes in fitness due to intra-cellular replication or losses due to cell death.

### Genetic relatedness among plasmids

So far we have focused on a partitioning of natural selection into among-group and within-group components, however alternative partitions exist, providing equivalent summaries of the total effects of natural selection [28, 29]. At this point, we turn briefly to a kin-selection partition of natural selection into individual (direct, *C*) and neighbourhood (indirect, *B*) components, with the indirect component weighted by a relatedness coefficient, *R*, such that a social trait is favoured if *RB* > *C* [2, 28, 29]. Here we focus solely on the the relatedness coefficient *R*, as a compact measure of population structuring, which can be translated for group traits into the proportion of total genetic variance among groups [29, 30]. The literature on the evolution of policing has focused extensively on the importance of relatedness as key determinant of selection for policing, with policing predicted to be favoured as a mechanism to maintain cooperative traits under conditions of low relatedness [3–6].

The relatedness coefficient with respect to a plasmid trait *z* (which can represent either of the plasmid parameters *β, κ,* or *α*) can be calculated for all plasmids in the population as the regression coefficient of the mean value of the trait 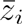 within each host *i* on the trait value *zij* of individual plasmids *j*, according to:

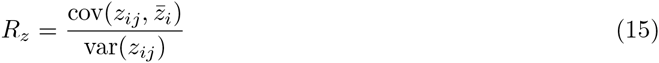

where the host-mean 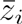 is calculated for each focal plasmid by excluding the parameter value of the focal plasmid itself (others-only relatedness) [30].

## Results

Our host/plasmid simulations consider the evolution of the plasmid replication parameters *β*, *κ* and *α*, with *ϕ* = 0.1, *γ* = 0.003, *λ* = 3, *ω*_0_ = 0.05, yielding an optimal copy number 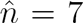 (see Equation 2). Each simulation starts with an inactive CNC mechanism among plasmids, i.e. initially plasmids neither produce replication inhibitors (*α* = 0) nor respond to their presence (*κ* = 0) and the plasmid parameters are allowed to evolve with a mutation probability *µ* = 0.005 per plasmid replication.

### Multi-level selection on plasmid replication

Figure 1 demonstrates the coevolution of the plasmid replication parameters over the course of a typical simulation leading to the emergence of stable plasmid replication control. The evolution of the plasmid-coded policing mechanism is driven, on one hand, by selection within hosts which operates directly on plasmid parameters so as to increase the intra-cellular replication rate of individual plasmids, and on the other hand by selection among hosts for higher rates of host reproduction. We discuss the influences of each of these levels of selection in turn.

**Figure 1.**
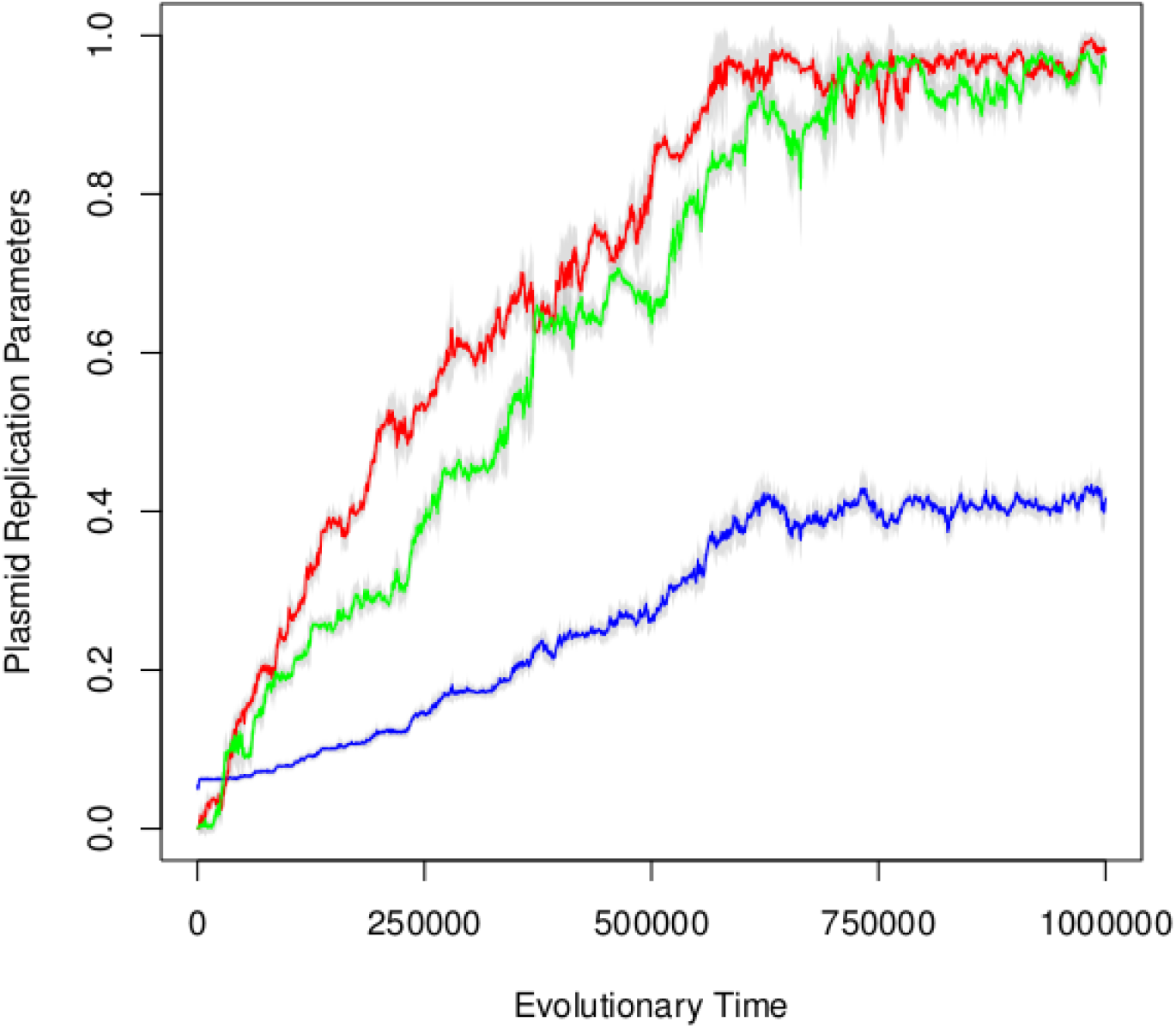
The coevolution of plasmid replication parameters. The evolutionary dynamics of the average plasmid basal replication rate 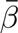 (blue), the average rate 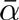 of production of policing resources (green), and the average obedience 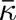 of plasmids to policing (red). The grey regions above and below the curves represent the standard deviation of each plasmid parameter in the population.

### Within-host selection

Figure 2 demonstrates that, on average, within-host selection favors cis-selfishness, i.e. higher baseline reproduction rates *β* (Figure 2A, red curve) and lower obedience *κ* to policing (Figure 2B, red curve). This is due to the fact that increased cis-selfishness translates into more frequent intra-cellular replication events, either because of higher intrinsic replication rates or because of lower obedience to policing, thus yielding a positive correlation with individual plasmid fitness.

**Figure 2.**
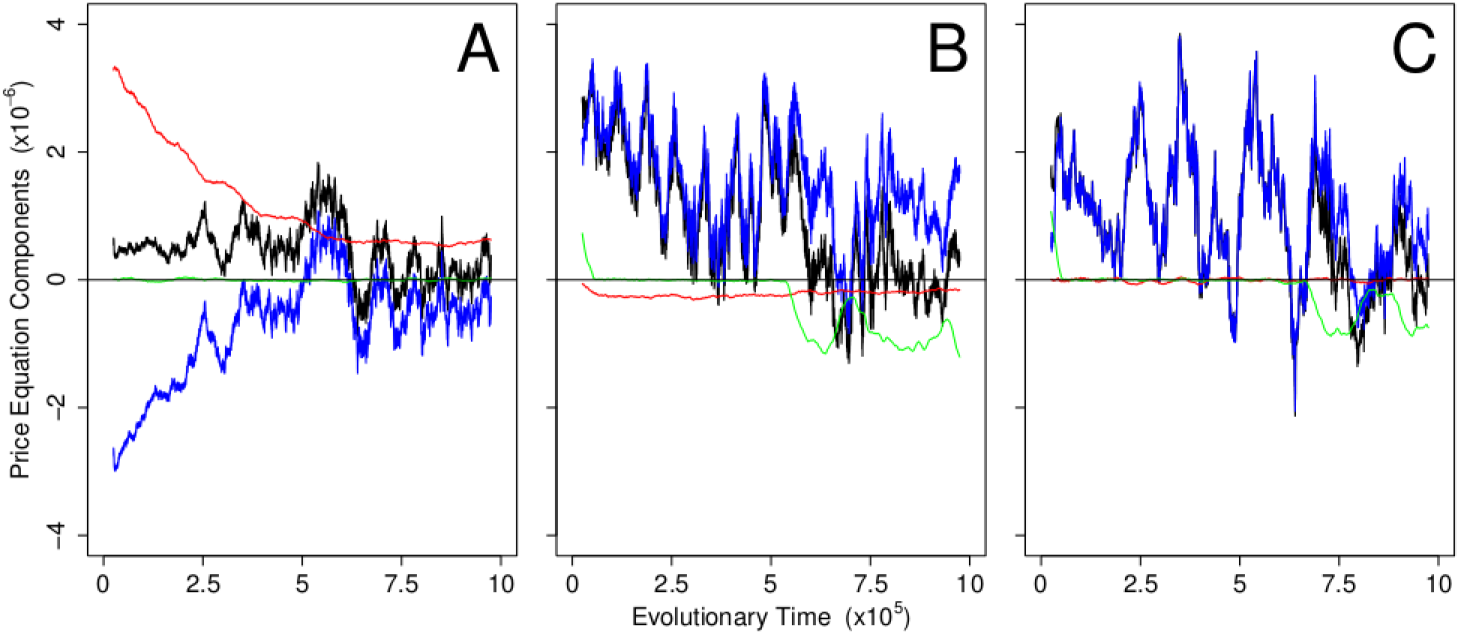
Partitioning selection on plasmids. The dynamics of the Price equation components (moving averages are shown) of the three plasmid replication parameters *β* (panel A), *κ* (panel B) and *α* (panel C) for the simulation shown in Figure 1. The effects of total selection on each plasmid parameter (black) are decomposed into between-host selection (blue), within-host selection (red) and the transmission bias term (green), according to Equation 11.

The trans-acting nature of the policing trait suggests that an increase in the production rate *α* of policing resources of an individual plasmid will influence all plasmids in that host through the same aggregate term 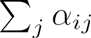 (see Equation 3). As a result, within-host selection on the policing trait is relatively neutral (Figure 2C, red curve).

### Between-host selection

Conversely, at the level of hosts, selection opposes plasmid cis-selfishness on average, by favoring lower basal replication rates *β* (Figure 2A, blue curve), and higher obedience *κ* to policing (Figure 2B, blue curve). Between-host selection also favors the production of policing resources (Figure 2C, blue curve), due to their positive impact on the stability of plasmid replication. However, selection at the level of hosts can occasionally favor plasmid cis-selfishness, as demonstrated by time frames in Figure 2 in which between-host selection (blue lines) is positive for *β* or negative for *κ*, *α*. This behaviour is transient and constitutes a response that alleviates the excessive inhibition that causes plasmids to under-replicate as a result of preceding increases in obedience *κ* and/or policing *α*.

Furthermore, Figure 2A (red line) shows that the progressive co-evolution of obedience and policing has a negative effect on within-host selection for intrinsic plasmid selfishness, as the gradual strengthening of the policing mechanism reduces the value of opportunistic plasmid behaviour in the intra-cellular replication pool. In other words, as policing becomes more dominant, the effect of *β*-increasing mutations on the replication rate of individual plasmids has a diminishing intensity.

Figure 2 also reveals the circumstances under which the transmission bias has an active role in shaping the evolutionary dynamics of the plasmid replication parameters. Specifically, the transmission bias terms for *κ* and *α* (green lines in Figures 2B and 2C) are positive at the start of the simulation, reflecting the fact that the parameters are very close to their minimum allowed value, and negative at the equilibrium phase, as *κ* and *α* are fluctuating just below their maximum allowed value (see also Figure 1). In the latter case, mutations will be predominantly negative (see Equation 4), thus counter-balancing the net positive selection for more policing and obedience (see corresponding black lines in Figures 2B and 2C). The existence of these parameter value limits, and in particular the upper limit, act as a constraint on the ratcheting effect in which the succession of selfish (higher *β*) and cooperative (higher *κ* and/or *α*) mutations would continue indefinitely, limited only by the costs of producing the factors involved in plasmid replication (initiators and inhibitors) [22].

### The evolution of policing

Figure 3 demonstrates that the emergence of policing and obedience among plasmids occurs against a background of consistently high relatedness (see Equation 15), suggesting that the plasmid population is highly homogeneous. In such cases, policing has been argued to be redundant as cooperation is predicted to be effectively maintained purely via kin selection [3]. Why, then, does policing evolve? An answer to this question is provided in Figure 4 (left) which shows that the evolution of the CNC mechanism has a positive effect on group performance even in highly related groups. This is demonstrated by the increasing division rate of hosts over time that results from the improving efficiency of control over the local population size, as measured by the host copy number.

**Figure 3.**
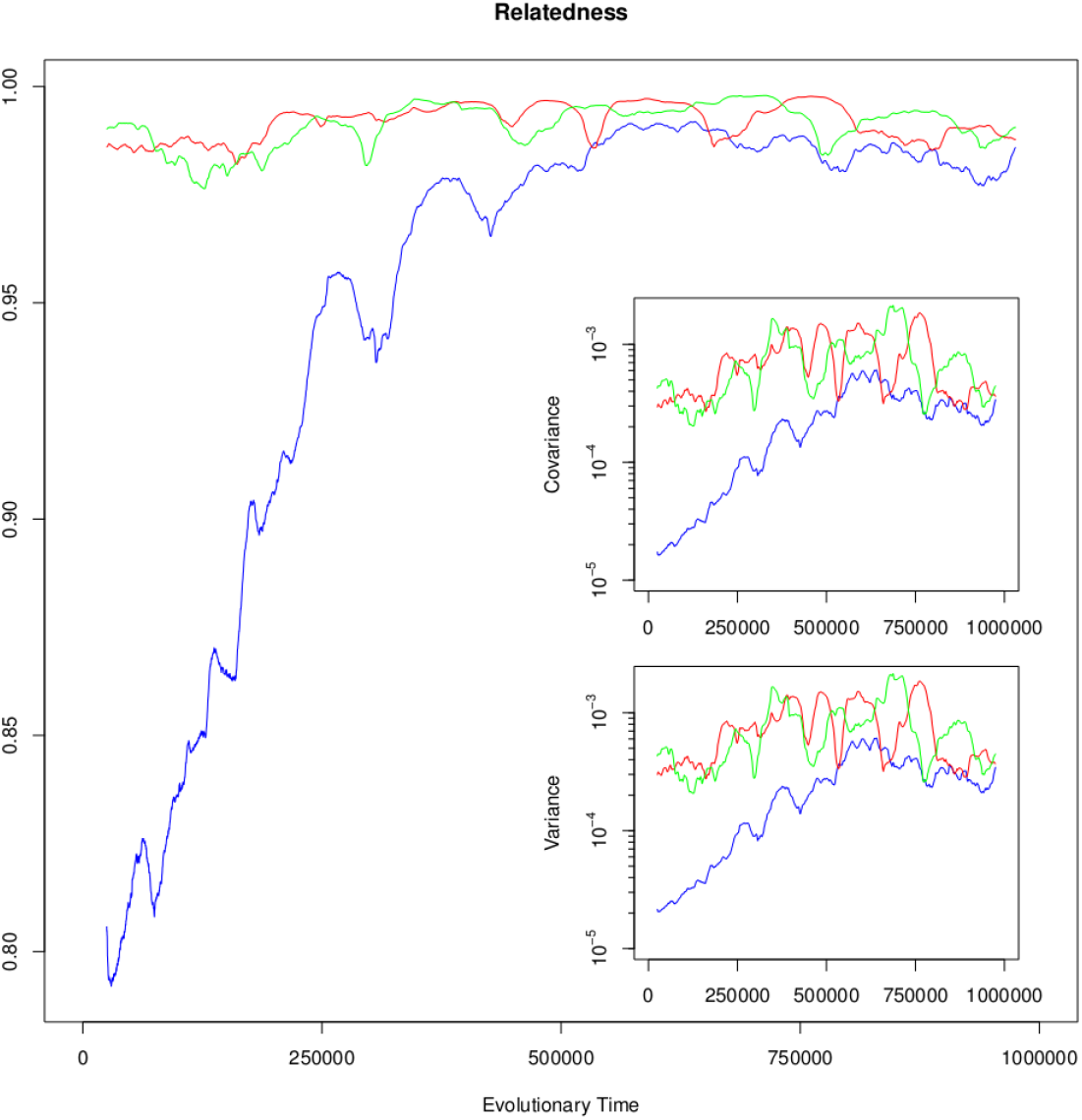
Policing evolves in the presence of high relatedness. The dynamics of the relatedness coefficient (moving averages are shown in main plot) for each of the plasmid parameters *β* (blue), *κ* (red) and *α* (green), calculated according to Equation 15 from the simulation shown in Figure 1. The dynamics of the components of the relatedness coefficient, 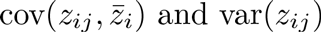, are also provided (inset plots).

**Figure 4.**
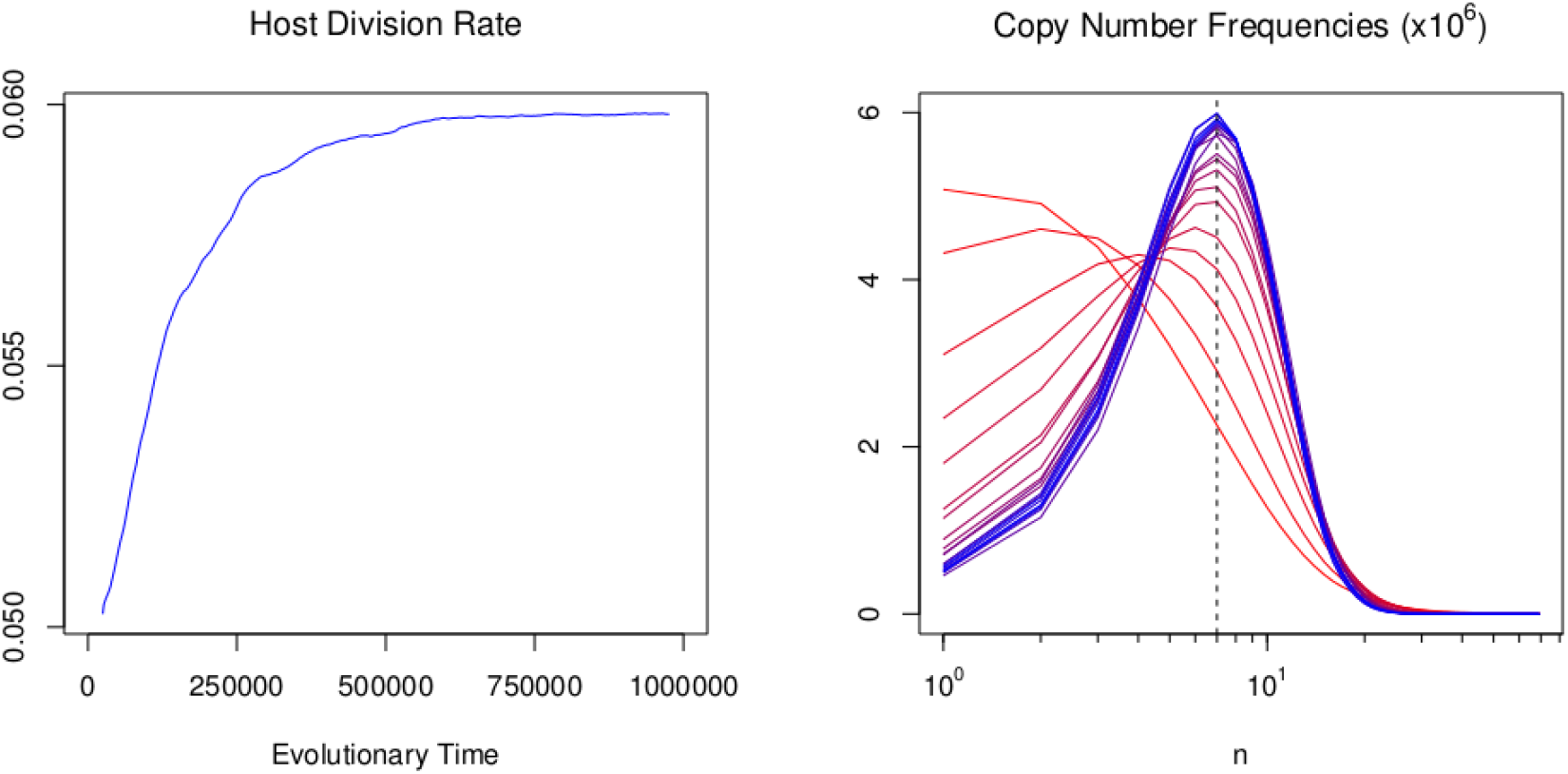
Policing improves group performance. The dynamics of the host division rate in the population (left) and the distributions of copy numbers in the population (right) calculated during consecutive time frames of 50000 steps from the simulation shown in Figure 1, starting from low levels of policing (red) up to a fully evolved CNC mechanism (blue). The dashed vertical line represents the copy number 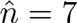 that is optimal for host growth (see also Equation 2).

The relationships between group size and performance can be explored by further decomposing the evolutionary pressures on the plasmid replication parameters and comparing the situations of low and high policing corresponding to different phases of the simulation. In particular, we can attribute selection to specific types of plasmid-related events such as intra-cellular replication and loss due to host death (see Equations 13 and 14) and ask the question how are these related to the evolution of the plasmid replication parameters. Figure 5 demonstrates that, when policing is absent or sufficently low (first phase of the simulation, up to 200K steps), selection at the level of hosts attributed to intra-cellular replication (blue lines) acts in favor of cis-selfishness and against policing. This obervation appears to be at odds with our results so far, namely that selection at the level of hosts should favor, rather than oppose, policing. However, we have to keep in mind that between-host selection does not operate directly on the plasmid replication parameters but, rather, on the distribution of copy numbers in the population [20] with the objective of maximizing host growth (see Equation 1). More specifically, between-host selection acts so as to align the mean of the population-wide copy number distribution to the optimal copy number 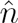, and to decrease the distribution’s variance in order to mitigate the intrinsic instabilities of plasmid replication [22]. These effects are demonstrated in Figure 4 (right), which also reveals that at the regime of no or low policing (red lines), the mean of the copy number distribution is lower than the copy number 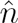 that is optimal for host growth. This explains why, at that regime, selection at the level of hosts attributed to intra-cellular replication is in favor of cis-selfishness and against policing: so that the mean copy number can increase and approach the value 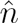 that is optimal for host growth.

**Figure 5.**
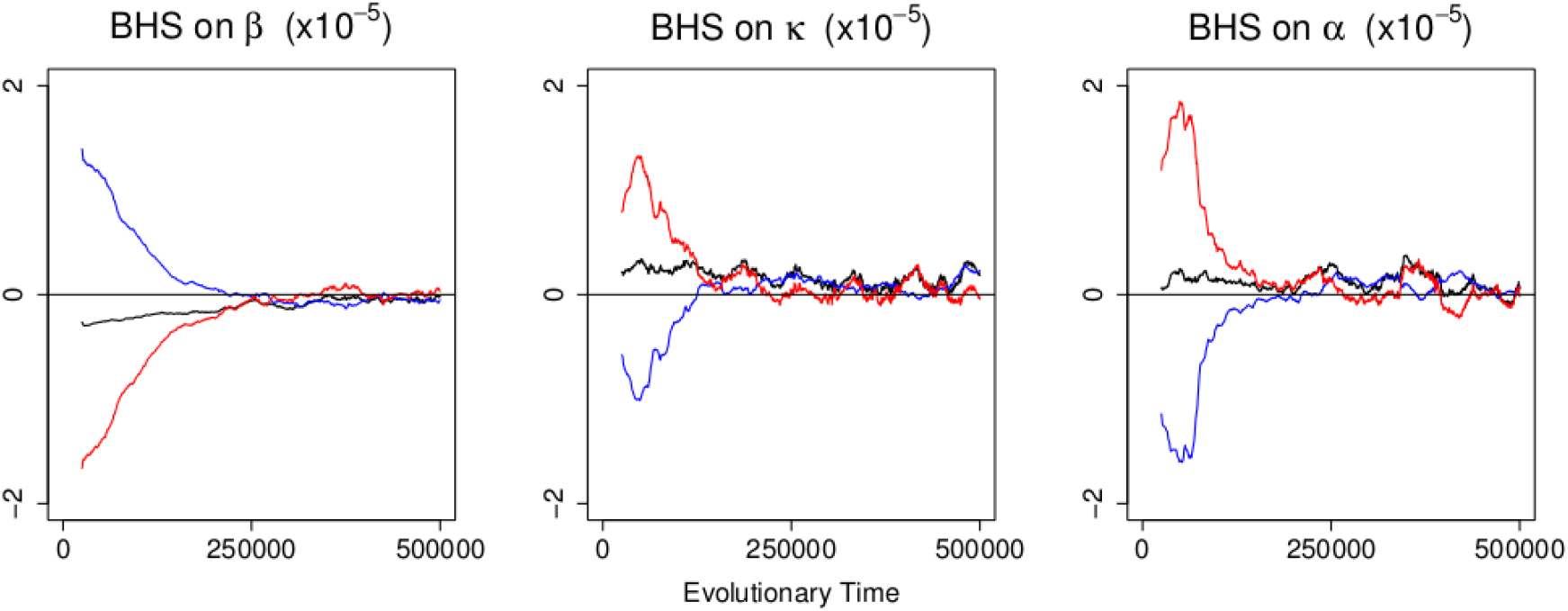
Decomposing between-host selection on the plasmid replication parameters. The dynamics of between-host selection (BHS) on the plasmid replication parameters (black) and their decomposition into a portion attributed to intra-cellular replication (blue) and a portion attributed to plasmid loss due to host death (red) according to Equation 13, calculated from the first half of the simulation shown in Figure 1. The behaviour at the regime of low policing (up to step 200K) is markedly different; specifically, the intra-cellular replication component is in favor of cis-selfishness and against policing, while the plasmid loss component is against cis-selfishness and in favor of policing.

However at the same time, this drive for higher copy numbers at the low policing regime exacerbates the inherent plasmid replication instabilities. The over-replication of selfish plasmids slows down host growth, thus reducing the propagation frequency of selfish plasmids through cell division. Such plasmids are more likely to suffer losses either due to the competition between hosts, which favors hosts that divide more frequently, or due to host death as a result of excessive over-replication. This effect is demonstrated in Figure 5 (red lines), which shows that, at the level of hosts, selection attributed to plasmid losses acts against cis-selfishness and in favor of policing, thus counter-balancing selection attributed to intra-cellular replication. Overall, the sum of these two evolutionary forces (see Equation 13) is in favor of policing as shown in Figure 5 (black lines). Furthermore, this conflict between the selective pressures at the level of hosts dies out as soon as obedience and policing evolve to sufficiently high levels so as to optimize the copy number distributions by eliminating the instabilities of plasmid replication, as shown in Figure 4 (right).

This analysis is not sensitive to the value of the optimal copy number 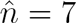 of the plasmid profile that was used in our simulations. Figure 6 shows the components of between-host selection for a simulation that uses a plasmid profile with a low optimal copy number 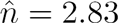 (with *ϕ* = 0.18, λ = 1, γ = 0.012). The dynamics demonstrate that, while the magnitude of the selective pressures is higher in the low optimal copy number simulations, their mode of operation remains fundamentally the same.

**Figure 6.**
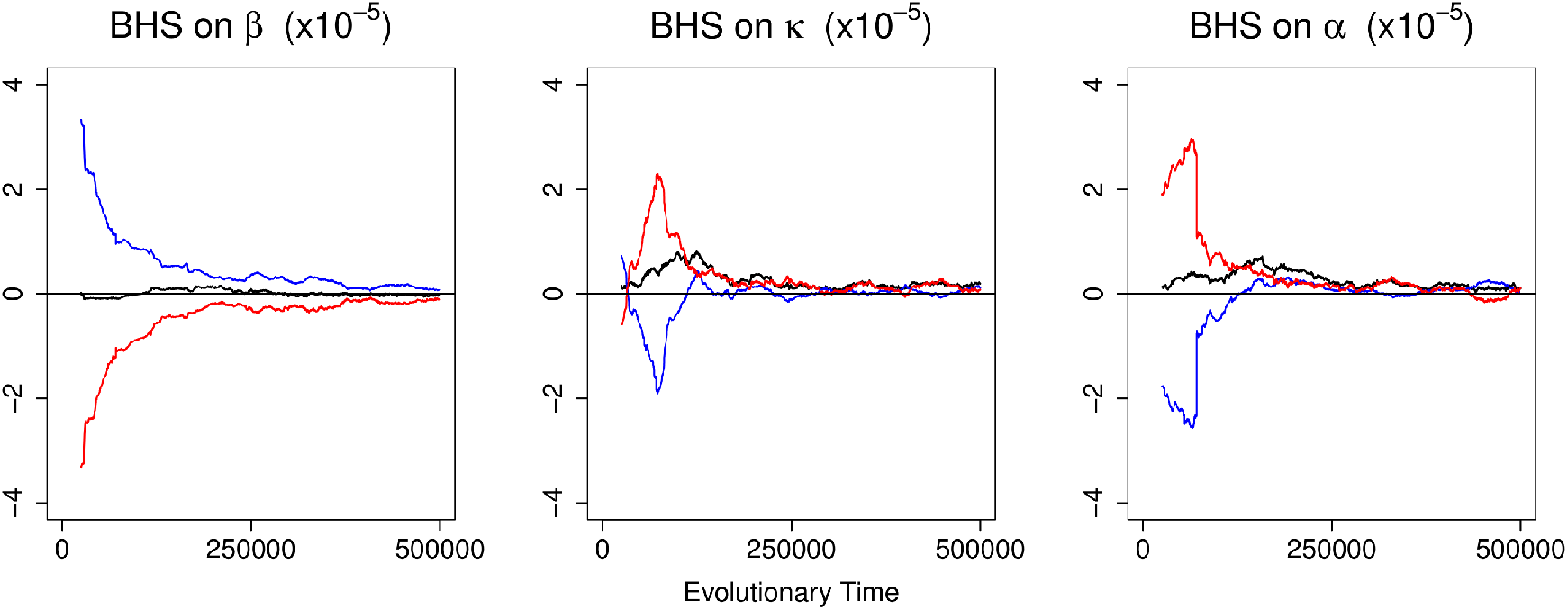
Decomposing between-host selection on the plasmid replication parameters for a low copy number plasmid. The dynamics of between-host selection (BHS) on the plasmid replication parameters (black) and their decomposition into a portion attributed to intra-cellular replication (blue) and a portion attributed to plasmid loss due to host death (red) according to Equation 13, calculated from the first half of the simulation. In contrast to Figure 5, the plasmid profile in use has a low optimal copy number 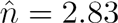 (with *ϕ* = 0.18, *γ* = 0.012 and *λ* = 1).

## Discussion

The emergence of stable reproductive control among plasmids is mediated by the coevolution of the plasmid replication parameters. Using the multi-level form of the Price Equation (see Equation 11), we were able to quantify and investigate the effects of selection on the plasmid CNC mechanism. The partitioning of selection into two components, namely between- and within- hosts, shed light on the underlying evolutionary pressures that shape the emergence and preservation of collective policing among plasmids. Specifically, we demonstrated the conflict between the levels of selection (Figure 2), by showing that selection within hosts favors plasmid cis-selfishness (i.e. higher *β* and lower *κ*), while, on the contrary, between-host selection opposes cis-selfishness and favors policing (higher *α*), due to the associated beneficial consequences on cellular growth.

The evolution of policing and the progressive intensification of the inhibitory effects of policing on intra-cellular replication was shown to reduce the value of cis-selfish plasmid mutations (Figure 2A – red line). In addition, it occured against a backdrop of consistently high genetic relatedness among plasmids (Figure 3). In contrast to the view that policing is redundant in the absence of within-group genetic conflict (high relatedness) [3], we found that policing evolves in a high-relatedness regime due to its positive effects on group efficiency and productivity resulting from better control over the local population size (Figure 4). We highlight that the reason for the discrepancy in predictions does not stem from the mode of analysis (kin versus group partitions of selection) but rather differences in the underlying evolutionary models.

By further decomposing between-host selection into the constituent components of plasmid fitness, namely the atomic events of intra-cellular replication and loss due to host death, we also showed how the plasmid replication instabilities at the low CNC regime drive the evolution of policing. Specifically, in the absence of policing (or when policing is low), between-host selection due to intra-cellular replication favors cis-selfishness and opposes policing (Figure 5 – blue lines). This is due to the shape of the population’s copy number distribution which, at low policing, yields a mean copy number lower than the optimal 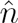 (Figure 4 – right), so that between-host selection attributed to intra-cellular replication favors higher replication rates and the growth of the mean copy number towards the optimal one. The drive towards higher replication rates exacerbates the plasmid replication instabilities and lowers the host division rate, hence plasmids that are more selfish will propagate less frequently in the population. This effect of plasmid losses due to host death generates selection at the level of hosts in favor of policing (Figure 5 – red lines) and constitutes the dominant factor in driving the evolution of plasmid replication control at the low policing regime.

Our approach serves as an illustrative example of the interpretative power of the Price equation. In itself, the Price equation has been argued to lack dynamic sufficiency in the sense that it only describes the change in the average character value and does not provide the means to generate the next state of the population [31]. However, the Price equation is a mathematical identity that can be neither dynamically sufficient nor dynamically insufficient as these characterizations are appropriate to models rather than algebraic identities [32, 33]. The use of the multi-level form of the Price equation (see Equation 11) provides insight into the interplay between the levels of selection by quantifying the effects of selection not only across hierarchical levels, but also across the different classes of events that determine plasmid fitness, namely intra-cellular replication and loss due to host death (see Equations 13 and 14). This allows for discerning and discussing the selective importance of different aspects of plasmid-related mechanisms and their relative contribution to the evolution of plasmid replication control.

